# Violation of the ultrastructural size principle in the dorsolateral prefrontal cortex underlies working memory impairment in the aged common marmoset (*Callithrix jacchus*)

**DOI:** 10.1101/2022.11.26.518060

**Authors:** Courtney Glavis-Bloom, Casey R. Vanderlip, Sammy Weiser Novak, Masaaki Kuwajima, Lyndsey Kirk, Kristen M. Harris, Uri Manor, John H. Reynolds

**Author notes:** These authors contributed equally (Courtney Glavis-Bloom, Casey R. Vanderlip). Corresponding author: Courtney Glavis-Bloom.

## Abstract

Working memory relies critically on the dorsolateral prefrontal cortex (dlPFC). Morphology and function of the dlPFC, and corresponding working memory performance, are affected early in the aging process. However, these effects are heterogeneous, with nearly half of aged individuals spared of working memory deficits. Translationally relevant model systems are critical for investigating the neurobiological drivers of this variability and identifying why some people experience age-related working memory impairment while others do not. The common marmoset (*Callithrix jacchus*) is advantageous as a model in which to investigate the biological underpinnings of aging because, as a nonhuman primate, marmosets have a clearly defined dlPFC facilitating investigations of prefrontal-dependent cognitive functions, including working memory, and their short (~10 year) lifespan facilitates longitudinal studies of aging. Here, we conduct the first investigation of synaptic ultrastructure in the dlPFC of the marmoset and investigate whether there are changes to synaptic ultrastructure that are unique to aging with and without working memory impairment. To do this, we characterized working memory capacity in a cohort of marmosets that collectively covered their short lifespan, and found age-related working memory impairment. We also found a remarkable degree of heterogeneity in performance, similar to that found in humans. Utilizing three dimensional reconstruction from serial section electron microscopy, we visualized structural correlates of synaptic efficacy including boutons, mitochondria, and synapses in layer III of the dlPFC of three marmosets: one young adult (YA), one aged cognitively unimpaired (AU), and one aged cognitively impaired (AI). We find that aged marmosets have fewer synapses in dlPFC than young, and this is due to selective vulnerability of small synapses. Next, we tested the hypothesis that violation of the ultrastructural size principle underlies age-related working memory impairment. The ultrastructural size principle states that synaptic efficacy relies on coordinated scaling of synaptic components (e.g., synapses, mitochondria) with presynaptic boutons. While synapses and mitochondria scaled proportionally and were strongly correlated with presynaptic boutons in the YA and AU marmosets, the ultrastructural characteristics of the AI marmoset were alarmingly different. We found that age-related working memory impairment was associated with disproportionately large synapses compared to presynaptic boutons, specifically in those with mitochondria. Remarkably, presynaptic mitochondria and these boutons were completely decorrelated. We posit that this decorrelation results in mismatched energy supply and demand, leading to impaired synaptic transmission. This is the first report of age-related synapse loss in the marmoset, and the first demonstration that violation of the ultrastructural size principle underlies age-related working memory impairment.

## INTRODUCTION

Working memory, the ability to maintain and manipulate information for short durations, critically depends on the dorsolateral prefrontal cortex (dlPFC) (Funahashi et al., 1989; Arnsten et al., 2012). Within the dlPFC, neurons in layer III fire repeatedly in the absence of selective sensory stimuli to hold information in working memory (Goldman-Rakic, 1995; Arnsten et al., 2012). The properties of this elevated firing is an area of active research and multiple models have been proposed (Constantinidis et al., 2018; Miller et al., 2018; Wang,2021). Importantly, these models posit that stimulus-selective changes in ongoing neural activity are necessary to support working memory. This activity is, by necessity, reliant on synaptic efficacy, the ability of synapses to efficiently transfer information between neurons. This efficacy is essential to the formation and maintenance of neural networks that underlie learning and memory (Kennedy, 2016). Therefore, intact memory is dependent on synaptic efficacy (Borczyk et al., 2021), and reduced synaptic efficacy results in decreased information transfer and memory impairment (Świetlik et al., 2019). Thus it is unsurprising that elevated firing that supports working memory is vulnerable to changes that compromise synaptic efficacy, such as those that occur with aging. Indeed, the dlPFC undergoes morphological and functional changes with age that correlate with working memory impairment (Morrison and Baxter, 2012; Upright and Baxter, 2021). For example, in macaque monkeys, aging and working memory impairment are associated with synapse loss (Hao et al., 2007; Peters et al., 2008). Selective loss of small synapses accounts for this loss (Dumitriu et al., 2010), and it is these synapses that are thought to be essential for the elevated firing in dlPFC that supports working memory (Kasai et al., 2003; Wang et al., 2011; Arnsten et al., 2012).

Elevated firing is metabolically demanding due, in part, to the high energy demands of synaptic transmission (Attwell and Laughlin, 2001), and this energy is primarily generated by presynaptic mitochondria (Rossi and Pekkurnaz, 2019; Pekkurnaz and Wang, 2022). Presynaptic mitochondria are a critical source of energy needed to support synaptic efficacy, and the demand for their energy is especially acute during elevated firing such as that which supports working memory. Therefore, it is unsurprising that the dlPFC has increased mitochondrial density compared to other cortical regions (Chandrasekaran et al., 1992; Arnsten et al., 2012).

As mitochondria age, they become impaired in their ability to provide energy, resulting in a reduced energy budget for the brain (Yao et al., 2010; Todorova and Blokland, 2017; Olesen et al., 2020). Aging is also associated with structural changes to mitochondria that are indicative of dysfunction. For example, in aged macaques with working memory impairment, there is increased prevalence of toroidal-shaped mitochondria in presynaptic boutons from layer III dlPFC. These boutons that contain toroidal-shaped mitochondria are associated with markers of decreased synaptic strength (Hara et al., 2014).

The ultrastructural size principle describes the fact that strong correlations between synaptic components (e.g., presynaptic mitochondria and synapses) and the size of the associated presynaptic bouton are critical for synaptic efficacy (Pierce and Mendell, 1993; Pierce and Lewin, 1994; Meyer et al., 2014). These correlations are upheld during periods of synaptic growth and remodeling (Meyer et al., 2014). Specifically, the elevated firing that supports working memory is defined by extended periods of synaptic activity during which the energy demands of synapses change rapidly. Under these conditions, it is especially important that each component of the synaptic complex varies in lockstep to meet the fluctuating energy demands of synapses and maintain effective synaptic transmission (Smith et al., 2016). The present study tests the hypothesis that violations of the ultrastructural size principle undermine synaptic efficacy, thereby contributing to synaptic dysfunction, and consequently, age-related working memory impairment. These violations would be observed as weak or absent correlations between the sizes of synaptic components in aged animals with working memory impairment compared to aged animals without cognitive impairment and young animals.

To test this hypothesis, we utilized three dimensional reconstruction from serial section electron microscopy (3DEM) to quantify structural correlates of synaptic strength in layer III dlPFC from three marmosets that varied in age and working memory capacity. Using this approach, we quantified synaptic components including presynaptic boutons, presynaptic mitochondria, and synapses, and investigated their relationship with each other. We identified a pronounced violation of the ultrastructural size principle that is specific to the aged marmoset with working memory impairment.

## METHODS

### Subjects

Two aged common marmosets (*Callithrix jacchus*, 9.3 year-old male, 9.6 year-old female), previously assessed on a working memory task (Glavis-Bloom et al., 2022), and one young control (1.3 years-old female), were used for this study. All experiments were conducted in compliance with the Institutional Animal Care and Use Committee of the Salk Institute for Biological Studies and conformed to NIH guidelines.

### Cognitive testing

Working memory ability of the aged marmosets was assessed using the Delayed Recognition Span Task (DRST; Figure 1A), as described in detail previously (Glavis-Bloom et al., 2022). Briefly, the DRST was initiated by the marmoset touching a blue square stimulus in the center of the screen. Then, a single black and white stimulus, selected randomly without replacement from a bank of 400, was displayed on the screen in one of nine possible locations, also selected randomly. When the marmoset touched this first stimulus, a small liquid reward was dispensed. Following a two-second delay, during which the screen was blank, a forced choice presented two alternatives: the original stimulus appearing in its original location, and a second, novel stimulus, appearing in a different location. If the marmoset selected the novel object (non-match-to-sample), a correct response was logged, a liquid reward was dispensed, and another two-second delay ensued. Following the delay, the first two stimuli appeared in their original locations, and a third, novel stimulus, also appeared, in a pseudo randomly chosen location. The marmoset was again rewarded for choosing the novel stimulus. Novel stimuli were added after subsequent delays until the trial was terminated in one of three ways: 1) the marmoset made nine correct selections in a row; 2) the marmoset failed to make a selection within 12 seconds (i.e., omitted); 3) the marmoset made an incorrect response (i.e., selected a non-novel stimulus).

**Figure 1.**
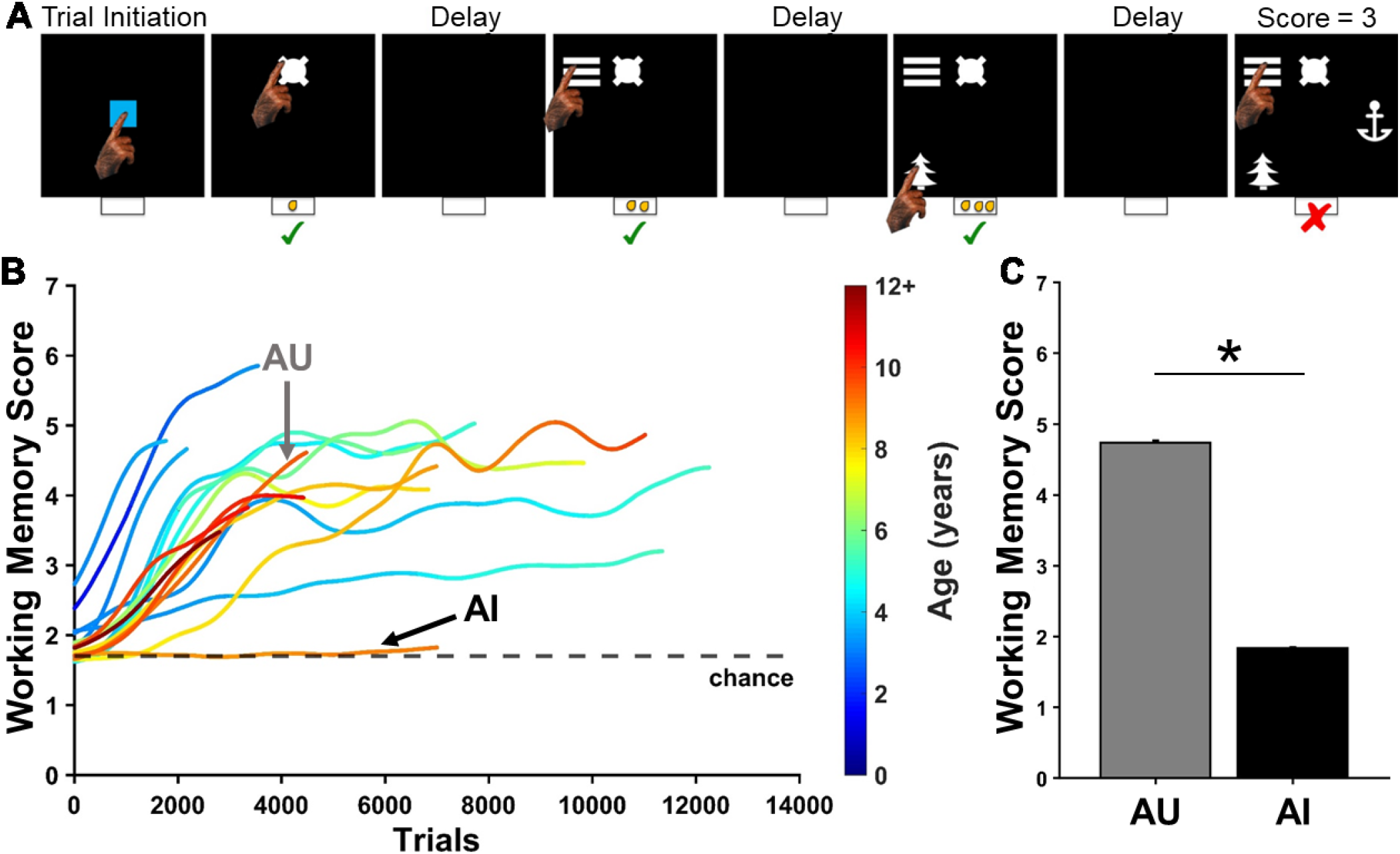
The Delayed Recognition Span Task (DRST) measures working memory. A) Depiction of one DRST trial. Green checks indicate correct responses. Red X indicates an error. B) Previously published learning curves for each of the 16 marmosets trained to perform the DRST. Chance performance is indicated by the black dashed line. Colors of learning curves indicate the age of the marmoset, with color changing within each curve as the animal aged over the course of the experiment. Working memory score is calculated as the average number of correct responses per trial. C) Average working memory score calculated over the last 500 DRST trials completed for each of two marmosets that performed differently on the DRST (mean ± SEM). *p < 0.05.

### Tissue collection

Even brief hypoxic episodes affect the structural integrity of synaptic ultrastructure which then precludes accurate ultrastructural measurements (Tao-Cheng et al., 2007). Therefore, we adapted a rodent hypoxia-resistant transcardiac perfusion euthanasia protocol (Kuwajima et al., 2013) for use in marmosets.

Alfaxalone (12 mg/kg, i.m.) and diazepam (2 mg/kg, i.m.) were administered to induce anesthesia, and atropine (0.02 mg/kg, i.m.) was administered to reduce saliva. Marmosets were deeply anesthetized with isoflurane (3-5%) delivered with oxygen (1-1.5L/min) via a mask over their nose and mouth, and then intubated and ventilated (2.75cc tidal volume, 60 breaths/min) to ensure oxygenation throughout the procedure. Once a deep surgical plane of anesthesia was reached, the thoracic cavity was opened with a lateral incision under the rib cage and a midline incision over the sternum. The pleura was bluntly dissected to expose the heart and lungs, and retractors were placed to maintain access to the thoracic cavity. The right atrium was clipped with iris scissors, and a blunt 18 gauge needle with warmed (37°C), oxygenated Krebs-Ringer Carbicarb (KRC) buffer flowing was inserted into the left ventricle. After three seconds, the marmosets were perfused with 2% formaldehyde and 2.5% glutaraldehyde in 0.1M cacodylate buffer (pH: 7.35). After one hour of perfusion, the brain was harvested, postfixed in the same fixation solution at 4°C for 48 hours, and then processed for electron microscopy.

### Electron microscopy

Materials were sourced from Electron Microscopy Sciences unless noted otherwise. Brains were processed into coronal sections of 100μm thickness using a vibrating microtome (Leica VT1000S, Leica Microsystems, Wetzlar, Germany). Sections that included the dorsolateral prefrontal cortex (dlPFC) area 46 (Paxinos et al.,2012) were selected for further processing. Sections were microdissected perpendicular to the cortical surface to produce samples that contained both the cortical surface and regions of interest (ROIs) in layer III, approximately 0.5mm tall and 3mm wide. Samples were transferred into glass vials and rinsed with ice cold 0.1M sodium cacodylate with 3mM calcium chloride for 15 minutes three times before staining and post-fixation with reduced osmium (1.5% osmium tetroxide, 1% potassium ferrocyanide, 3mM calcium chloride, 0.1M sodium cacodylate) for 45 minutes in the dark at room temperature. Samples were rinsed three times with ice cold deionized water and left overnight in 1% aqueous uranyl acetate in the dark at 4°C. Samples were rinsed three times briefly in deionized water before serial dehydration in ascending concentrations of ice cold ethanol (20, 50, 70, 80, 90, 100, 100; 10 minutes per change). Samples were then infiltrated with Eponate 12 resin (Pelco; Product No. 18012) hard formulation, in a 1:1 mix with anhydrous ethanol overnight on a rotating mixer. The following day, the samples were further infiltrated with 2:1::resin:ethanol mix for 2 hours, then transferred from their vials into shallow polypropylene bottle caps filled with pure resin for two hours. Fresh resin was exchanged and the samples were further infiltrated for two hours before embedding in a silicone rubber mold and polymerization for 48 hours at 70°C.

Semi-thin (0.7-1μm) sections were collected using a diamond knife (DiATOME, Hatfield, PA) on an ultramicrotome (Leica UC7), with sections extending from the cortical surface to the deeper cortical layers. Layer III was identified by neuroanatomical landmarks on the semi-thin sections, including distance from the cortical surface and density of neuronal somas (Yuasa et al., 2010; Paxinos et al., 2012). For each sample, the blockface was carefully trimmed using a 90° diamond trim tool to a dimension of approximately 0.6 x 0.2mm in layer III, and a ribbon of approximately 200 serial sections (55-70 nm) was collected onto a silicon chip partially immersed in the knife boat and, as the water level was lowered using the peristaltic pump, the sections dried down onto the silicon substrate. The chip was mounted on an aluminum stub using a sticky carbon tab, and loaded into a scanning electron microscope (SEM; Sigma VP scanning electron microscope, Zeiss). Images were collected using a Gatan backscattered electron detector at a working distance of 7mm and an accelerating voltage of 3kV. Image maps of the series at variable resolutions were assembled using the extended raster scanning and control system (Eberle et al., 2015) (Atlas5, FIBICS, Ottawa) and layer III was identified.

An ROI from layer III was selected for imaging by overlaying a numbered grid with 50μm x 50μm spacing, and randomly selecting a number from the grid. These ROIs were evaluated every 10 sections through the ribbon at an intermediate resolution to ensure that the series would be devoid of somas and blood vessels to ensure equivalent analysis of neuropil across samples. The ROIs were imaged through 41 consecutive serial sections with a pixel size of 2nm. The image sequence was contrast normalized using the scikit-image implementation of CLAHE (Walt et al., 2014) and rigidly aligned using TrakEM2 in Fiji (Cardona et al., 2012). These volumes were cropped to a minimum contiguous cube of roughly aligned data with no padding. Fine image stack alignment was accomplished using SWiFT-IR (https://ieeexplore.ieee.org/document/8010595) as deployed on 3DEM.org using the TACC compute resource Stampede 2 (Litvina et al., 2019), with final contiguous volumes formed of approximately 20μm x 20μm x 2.5μm.

### Image segmentation and reconstruction

The image stacks were imported into Volume Annotation and Segmentation Tool (VAST Lite) (Berger et al.,2018) for three-dimensional reconstruction of presynaptic boutons and their mitochondria, and the associated synapse surface area. We employed similar sampling methods to prior work in macaque monkeys (Hara et al.,2011, 2014, 2016; Fehr et al., 2022). The experimenter performing the segmentations was blind to the subject-identifying information of the images. The middle image of each stack (“reference image”) was selected, and a central region of dimensions 18μm x 9μm was identified. This 162μm^2^ analysis region was divided into four non-overlapping segmentation regions each 40.5μm^2^ (9μm x 4.5μm). Each of these four segmentation regions had two inclusion borders and two exclusion borders to standardize the volume of neuropil analyzed for each sample. Presynaptic boutons were identified on the reference region and were included if they were either fully inside a segmentation region, or partially crossed an inclusion border. These included boutons were segmented on the reference section and throughout their entirety through the image stack. As in prior work, presynaptic boutons were defined as containing three or more presynaptic vesicles on the reference section (Hara et al., 2014, 2016). Presynaptic mitochondria and synapses associated with these boutons were also segmented.

Three dimensional modeling software (Blender; www.blender.org) was used with the Geometry-preserving Adaptive Mesher plugin (GAMer) (Lee et al., 2019; Lee, 2020) to generate smoothed, biologically accurate contours (i.e., “meshes”). Bouton and mitochondria surface area and volume measurements were taken from the meshes, and the mitochondria were classified by their morphology (straight, curved, toroidal). In alignment with prior work, mitochondria that were bent less than 90 degrees were classified as straight, mitochondria that bent 90 degrees or more were classified as curved, and toroidal-shaped (sometimes referred to as donut-shaped in the literature),were classified as toroidal (Hara et al., 2014, 2016). Synapse surface area was quantified by measuring synapse lengths (i.e., apposition of the presynaptic active zone and postsynaptic density) from each image on which they appeared, and multiplying by the section thickness. Approximately 160 boutons were analyzed from each subject.

### Statistical analyses

Data were analyzed using MATLAB (Mathworks, Natick, MA). Shapiro-Wilk tests for normality revealed that the data were normally distributed, and so parametric statistical analyses were performed. Associations between two variables were assessed using Pearson correlations and Model II regressions. Strengths of the correlations were compared with Fisher Z tests. Two-way analysis of variance (ANOVA) tests were used to assess two factor interactions. One-way ANOVAs with Tukey HSD post-hoc tests were used to identify within-factor differences. P < 0.05 was considered significant.

## RESULTS

### Aging heterogeneously affects working memory capacity

Behavioral findings from a large group of marmosets, including the two aged monkeys used in this study, are described in detail elsewhere (Glavis-Bloom et al., 2022). Learning curves from each of these 16 animals are recapitulated in Figure 1B. Here, we focus on two aged marmosets that performed differently on the DRST; one successfully learned the rules of the DRST and reached a high level of performance (Aged Unimpaired; AU), while the other, similarly aged monkey, failed to learn the DRST, and continued to perform at chance levels of accuracy even after extensive experience on the task (Aged Impaired; AI). On the last 500 DRST trials each marmoset performed, the AI marmoset’s average Working Memory Score (number of items identified correctly on a given trial) was significantly lower than that of the AU marmoset (Figure 1C; Welch’s t-test: t(538) = 23.03, p = 3.45×10^-82^).

### Age-related working memory impairment is associated with reduced coordination of pre- and post-synaptic components

Synaptic efficacy is critically dependent on the correlated sizes of synaptic components including boutons, synapses, and mitochondria. This coordinated scaling is known as the ultrastructural size principle. Whether and how the ultrastructural size principle is upheld with age and in cognitive impairment, however, is unknown. To investigate the synaptic changes underlying this cognitive impairment, we used 3DEM to quantify synapses, boutons, and presynaptic mitochondria in layer III of the dlPFC from three marmosets: a young adult (YA), an aged cognitively unimpaired animal (AU), and an aged cognitively impaired animal (AI).

Prior observations in macaques and humans show that there is reduced synaptic density in layer III of the dlPFC with age, and this reduction is more severe in humans and non-human primates with age-related cognitive impairment (Peters et al., 2008; Dumitriu et al., 2010; Boros et al., 2017). Marmosets are shorter-lived than are macaques and humans, raising the question of whether this same pattern of synapse loss is observed with age and if it is more pronounced in aged marmosets exhibiting cognitive impairment. Consistent with prior work, we found a significant age-related decrease in synapse density (Figure 2A; one-way ANOVA: F(2) = 6.62, p = 0.006). However, synapse density was reduced in both AU and AI, and did not vary significantly with cognitive impairment (Tukey’s HSD: YA vs AU: p = 0.02; YA vs AI: p = 0.008; AU vs AI: p = 0.89). We then quantified bouton density and the number of synapses per bouton for each marmoset to evaluate whether the age-related reduction in synapse density was due to a loss of boutons, a reduction in the number of synapses per bouton, or a combination of the two. Similar to synapse density, we found an overall decrease in bouton density with age (Figure 2B; one-way ANOVA: F(2) = 8.05, p = 0.003) and no variation with working memory impairment (Tukey’s HSD: YA vs AU: p = 0.008; YA vs AI: p = 0.005; AU vs AI: p = 0.97). Furthermore, there was an equivalent number of synapses per bouton across animals (Figure 2C; one-way ANOVA: F(2) = 0.41, p = 0.67). Together, these results demonstrate synapse and bouton loss with age that does not differentiate with working memory impairment.

**Figure 2.**
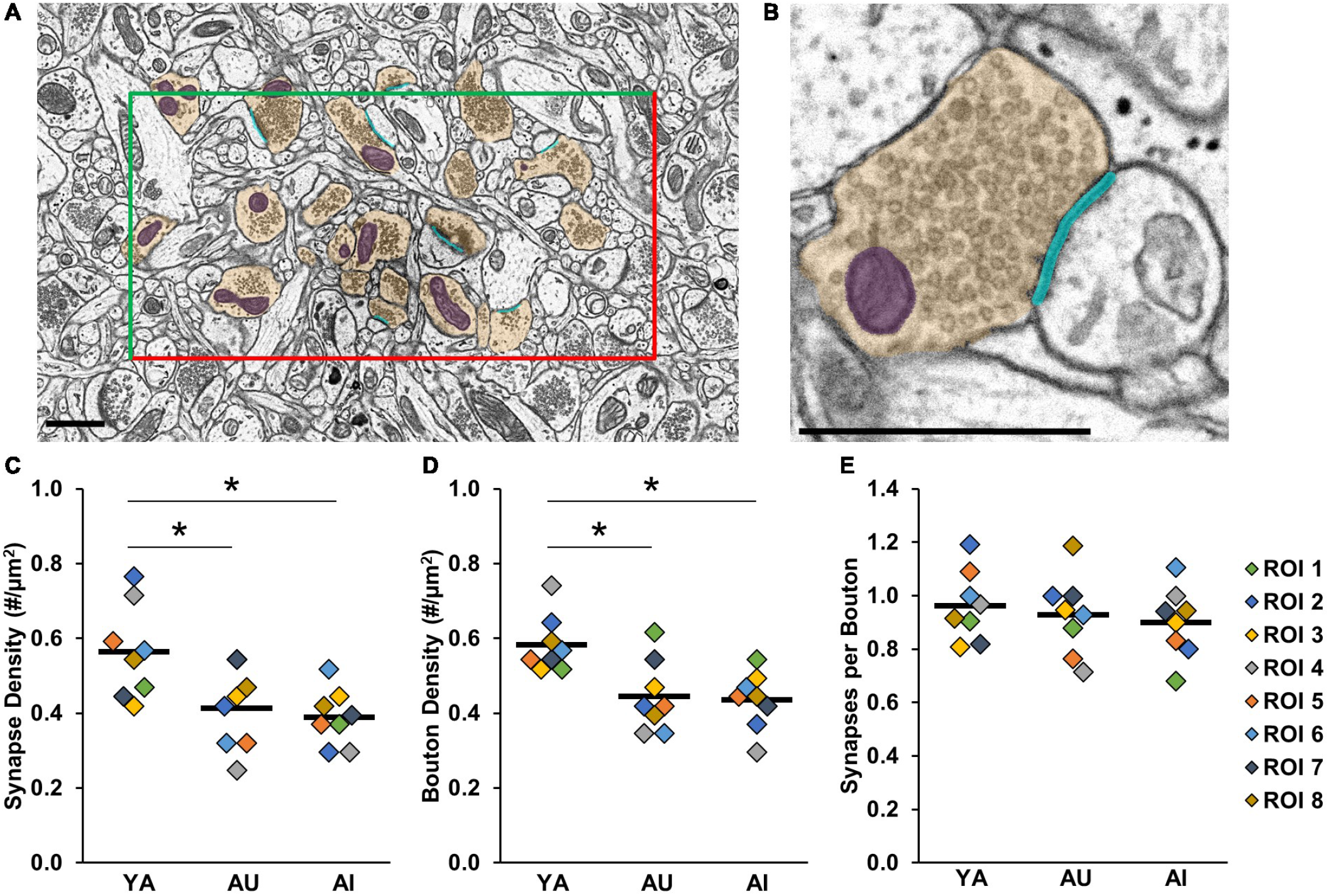
Synapse and bouton density are decreased with age regardless of working memory capacity. A) Reference section with one ROI outlined by two inclusion (green) and two exclusion (red) lines. Included presynaptic boutons are segmented in yellow, synapses in cyan, mitochondria in purple. Scale bar = 1μm. B) Zoomed in view of one synaptic complex. Colors and scale bar as in panel A. C) Synapse density D) Bouton density E) Equivalent number of synapses per bouton across animals. In panels C-E, colored diamonds represent measures taken in each of the ROIs. Horizontal black bars represent average across the ROIs. *p < 0.05.

This leaves open the possibility that, rather than being driven by synapse or bouton number, working memory impairments might stem instead from a dysfunction of the synapse itself. In particular, we hypothesized that a failure of coordination in the size of the components of the synaptic complex – violations of the ultrastructural size principle, might underlie cognitive impairment. To test this, we quantified synapse and bouton size in each of our marmosets (Figure 3), and then investigated whether violations of the ultrastructural size principle predicted cognitive performance.

**Figure 3.**
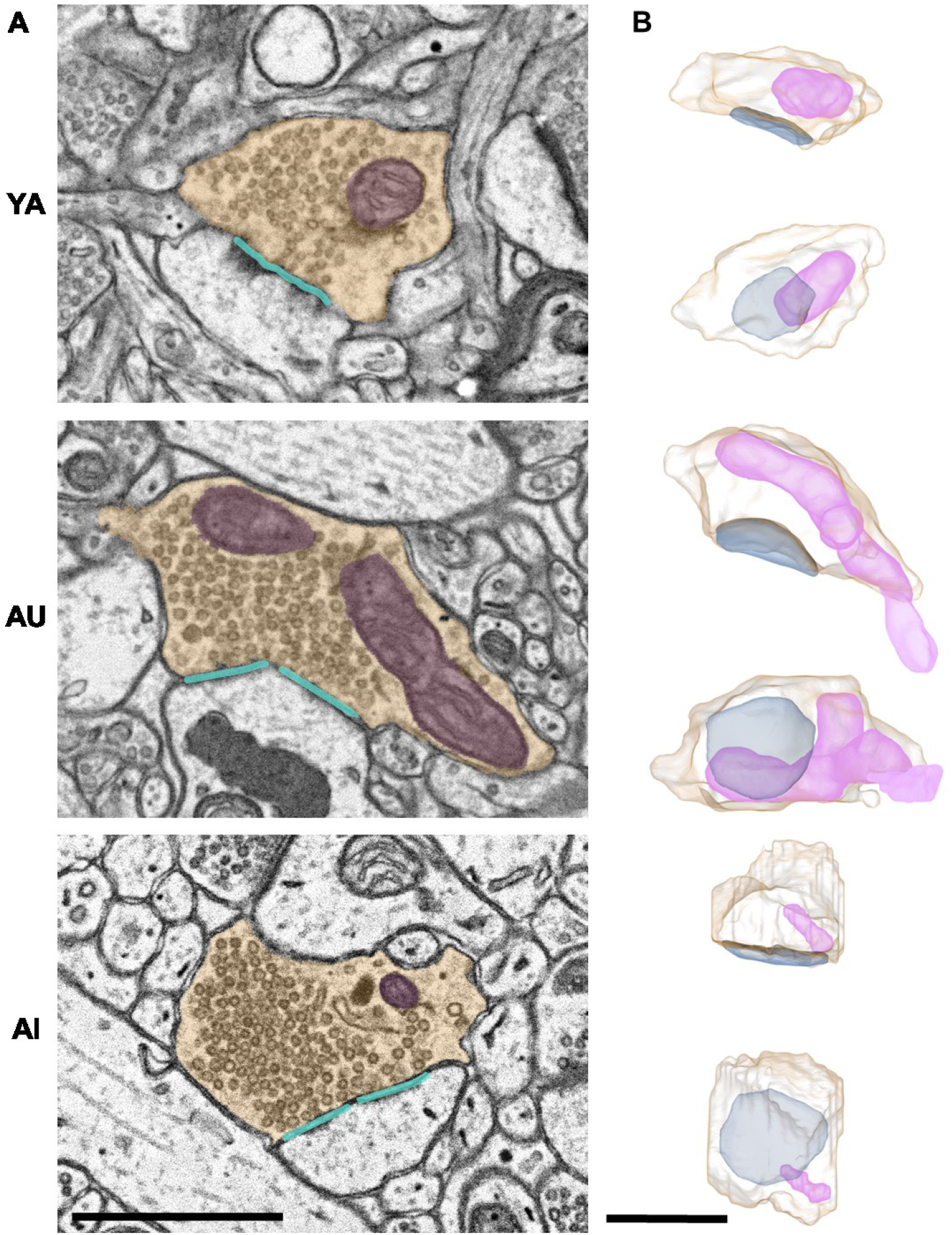
Synaptic complex segmentations and reconstructions. A) Two-dimensional segmentations, and B) paired orthogonal projections of representative synaptic complexes from the young adult (top row), aged unimpaired (middle row), and aged impaired (bottom row) marmosets. Presynaptic boutons are shown in yellow, mitochondria in purple, synapses in cyan. Scale bars are 1μm.

Consistent with prior microscopy studies from layer III dlPFC in macaques and humans, we observe an increased average synapse size with age that is exacerbated with cognitive impairment (Figure 4A; one-way ANOVA: F(2) = 31.23, p < 0.0001; Tukey’s HSD post-hoc: YA vs AU: p = 0.001; YA vs AI: p < 0.0001; AU vs AI: p = 0.004). Prior work has shown that the overall increase in average synapse size is due to a selective loss of small, plastic synapses with age, and further loss of these particular synapses with cognitive impairment (Dumitriu et al., 2010; Boros et al., 2017). Bouton size was similarly increased with age, however was not significantly changed with cognitive impairment (Figure 4B; one-way ANOVA: F(2) = 19.62, p < 0.0001; Tukey’s HSD: YA vs AU: p = 0.0003, YA vs AI: p = < 0.0001; AU vs AI: p = 0.54). Since the ultrastructural size principle states that synapse and bouton sizes are highly correlated, it is unexpected that synapse size varied significantly between AU and AI, and bouton size did not.

**Figure 4.**
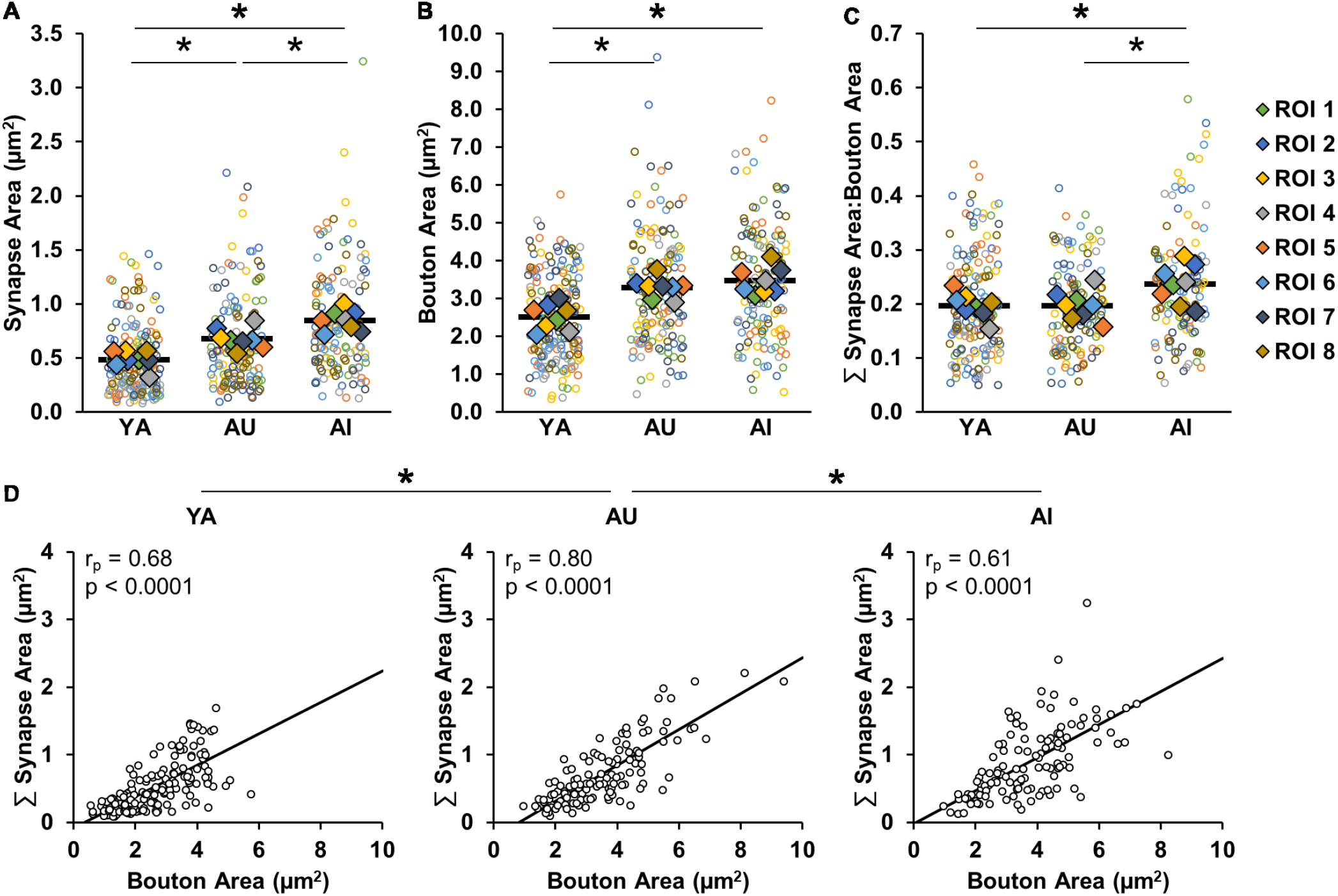
Relationship between synapse surface area and bouton surface area is altered with working memory impairment. A) Aged marmosets have larger synapse surface areas than young marmosets, with a further increase associated with working memory impairment. B) Bouton surface area was increased with age but there was no further increase associated with working memory impairment. C) Age-related working memory impairment is uniquely associated with an increased ratio of synapse surface area to bouton surface area. D) There are strong correlations between synapse size and bouton size in all conditions, with a significantly weaker correlation in the AI compared to the AU marmoset. In all panels, open circles show each individual datapoint. In A, B, & C, colored diamonds represent the average of measures taken in each of the ROIs, and horizontal black bars represent average across the ROIs. * p < 0.05.

To assess whether the reduced coordination between synapse and bouton size is specific for the AI marmoset, or whether it is ubiquitous with aging but exacerbated with age-related working memory impairment, we compared the ratios and correlations of synapse and bouton surface area across animals. The ratio of summed synapse surface area to bouton surface area was consistent between YA and AU, whereas it was larger in the AI marmoset (Figure 4C; one-way ANOVA: F(2) = 4.87, p = 0.02; Tukey’s HSD: YA vs AU: p = 1.00; YA vs AI: p = 0.03; AU vs AI: p = 0.03). We found strong correlations in all subjects (Figure 4D; YA: r_p_ = 0.68, p < 0.0001; AU: r_p_ = 0.80, p < 0.0001; AI: r_p_ = 0.61, p < 0.0001), and that the correlation was significantly weaker in AI compared to AU (Fisher Z transformation: YA vs AU: p = 0.024; YA vs AI: p = 0.32; AU vs AI: p = 0.003). In violation of the ultrastructural size principle, weaker correlation between boutons and synapses, along with disproportional scaling between these two components may reflect synaptic dysfunction that underlies the working memory impairment in AI.

### Synaptic boutons that contain mitochondria are selectively vulnerable to synaptic dysfunction in an aged marmoset with working memory impairment

We next asked whether the presence of presynaptic mitochondria affected the synapse to bouton ratio. First, we found no changes in the frequency of mitochondria containing boutons in any of the conditions (One-way ANOVA: F(2) = 0.75, p = 0.49). This opens the possibility that synapse and bouton surface area may vary as a function of the presence or absence of presynaptic mitochondria, and also with age and working memory impairment.

In regards to bouton size, we found that, across all animals, boutons with presynaptic mitochondria are larger than boutons without presynaptic mitochondria (Figure 5A; Two-way ANOVA: mitochondria presence F(1) = 160.31, p = 1.42×10^-15^; Tukey’s HSD: p = 1.63×10^-10^). Analyzed separately, however, there were differences between the animals both for boutons with mitochondria and boutons without mitochondria (Two-way ANOVA: animal F(2) =16.29, p =6.69×10^-6^; interaction (mitochondria presence x animal) F(2) = 8.49, p = 0.0008). For boutons without mitochondria, those in the AI marmoset were significantly larger than in YA and AU (Tukey’s HSD; YA vs AU: p = 0.35, YA vs AI: p = 0.0008, AU vs AI: p = 0.02). For boutons with mitochondria, boutons were significantly larger in both aged marmosets compared to young, however AI marmoset boutons with mitochondria were significantly smaller than those in AU (Tukey’s HSD; YA vs AU: p = 3.68×10^-5^, YA vs AI: p = 0.02, AU vs AI: p = 0.03).

**Figure 5.**
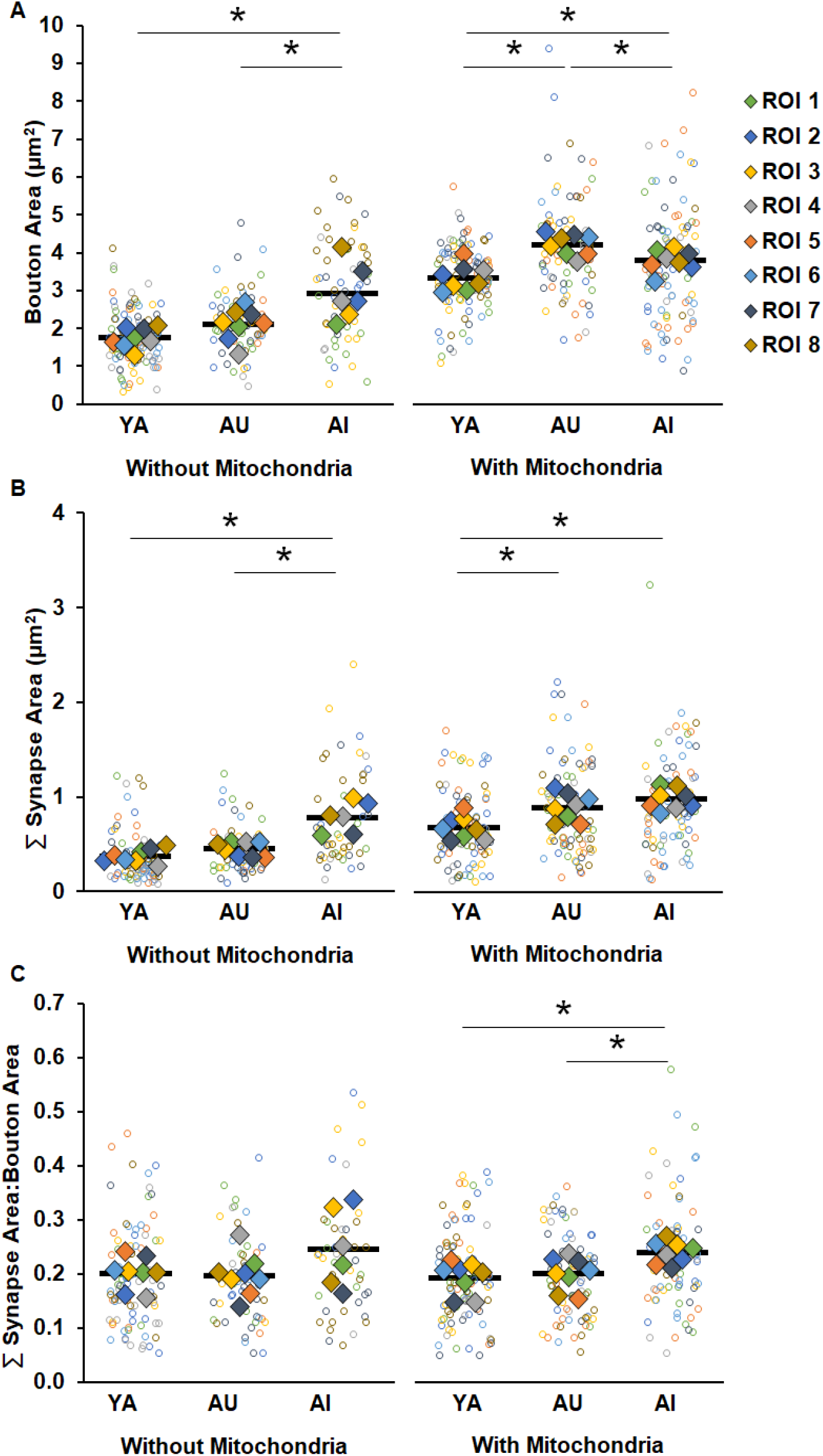
Boutons and synapses with and without mitochondria. A) Boutons without presynaptic mitochondria were larger in AI compared to YA and AU. Boutons with presynaptic mitochondria were larger in both aged marmosets than the young, with further enlargement in AU compared to AI. B) Synapses without associated presynaptic mitochondria were larger in AI compared to YA and AU, while synapses with associated presynaptic mitochondria were increased in size with age regardless of cognitive status. C) Ratio of synapse size to bouton size is significantly larger in the AI marmoset compared to YA and AU only when presynaptic mitochondria were present. Open circles show each individual datapoint, colored diamonds represent the average of measures taken in each of the ROIs, and horizontal black bars represent average across the ROIs. * p < 0.05.

We applied these same analyses to assess the effects of mitochondrial presence on synapse size, and found a similar pattern of results (Figure 5B; Two-way ANOVA: mitochondria presence F(1) = 78.70, p < 0.0001; animal F(2) = 33.58, p < 0.0001; interaction (F(2) = 4.21, p = 0.02). Specifically, across all animals, synapses associated with presynaptic mitochondria were larger than those that were not (Tukey’s HSD: p < 0.0001). Also, synapses without associated presynaptic mitochondria were larger with cognitive impairment (Tukey’s HSD; YA vs AU: p = 0.72, YA vs AI: p < 0.0001, AU vs AI: p = 0.0001), whereas synapses associated with mitochondria were larger in both aged animals relative to YA, and did not vary with cognitive status (Tukey’s HSD; YA vs AU: p = 0.01, YA vs AI: p = 0.0001, AU vs AI: p = 0.72). Taken together, these results show that the presence or absence of presynaptic mitochondria has differential effects on structural correlates of synaptic strength, and these effects vary as a function of age and cognitive impairment.

To resolve the question of whether the presence or absence of presynaptic mitochondria renders associated boutons and synapses vulnerable to dysfunction, we calculated the ratio of synapse to bouton size separately for boutons without mitochondria and boutons with mitochondria (Figure 5C). We found a significant increase in the synapse to bouton size ratio of the AI marmoset compared to the YA and AU animals in the boutons containing mitochondria, but not in the boutons without mitochondria (One-way ANOVA: without mitochondria: F(2) = 2.18, p = 0.14, with mitochondria: F(2) = 6.77, p = 0.005; Tukey’s HSD: YA vs AU: p = 0.83; YA vs AI: p = 0.03; AU vs AI: p = 0.03). Therefore, mitochondria likely contribute to synaptic dysfunction in AI, but how this occurs is an open question.

### Decorrelation between presynaptic boutons and mitochondria underlie synaptic dysfunction in age-related working memory impairment

Synaptic transmission is metabolically demanding and this energy is primarily supplied by mitochondria (Attwell and Laughlin, 2001) that are essential for synaptic efficacy and stability (Smith et al., 2016; Lees et al., 2020). While the percentage of boutons that contained mitochondria did not vary across animals (Figure 5A; one-way ANOVA: F(2) = 0.75, p = 0.49), there are several mechanisms by which mitochondria could underlie synaptic dysfunction, including: 1) low mitochondrial density, 2) dysmorphic mitochondria, 3) reduced mitochondrial size, and 4) improper scaling of mitochondria and bouton size. We investigate each of these in-turn, below.

We first quantified the number of mitochondria per bouton for each animal, and found that the average did not vary across animals (Figure 6A: one-way ANOVA: F(2) = 1.72, p = 0.20).

**Figure 6.**
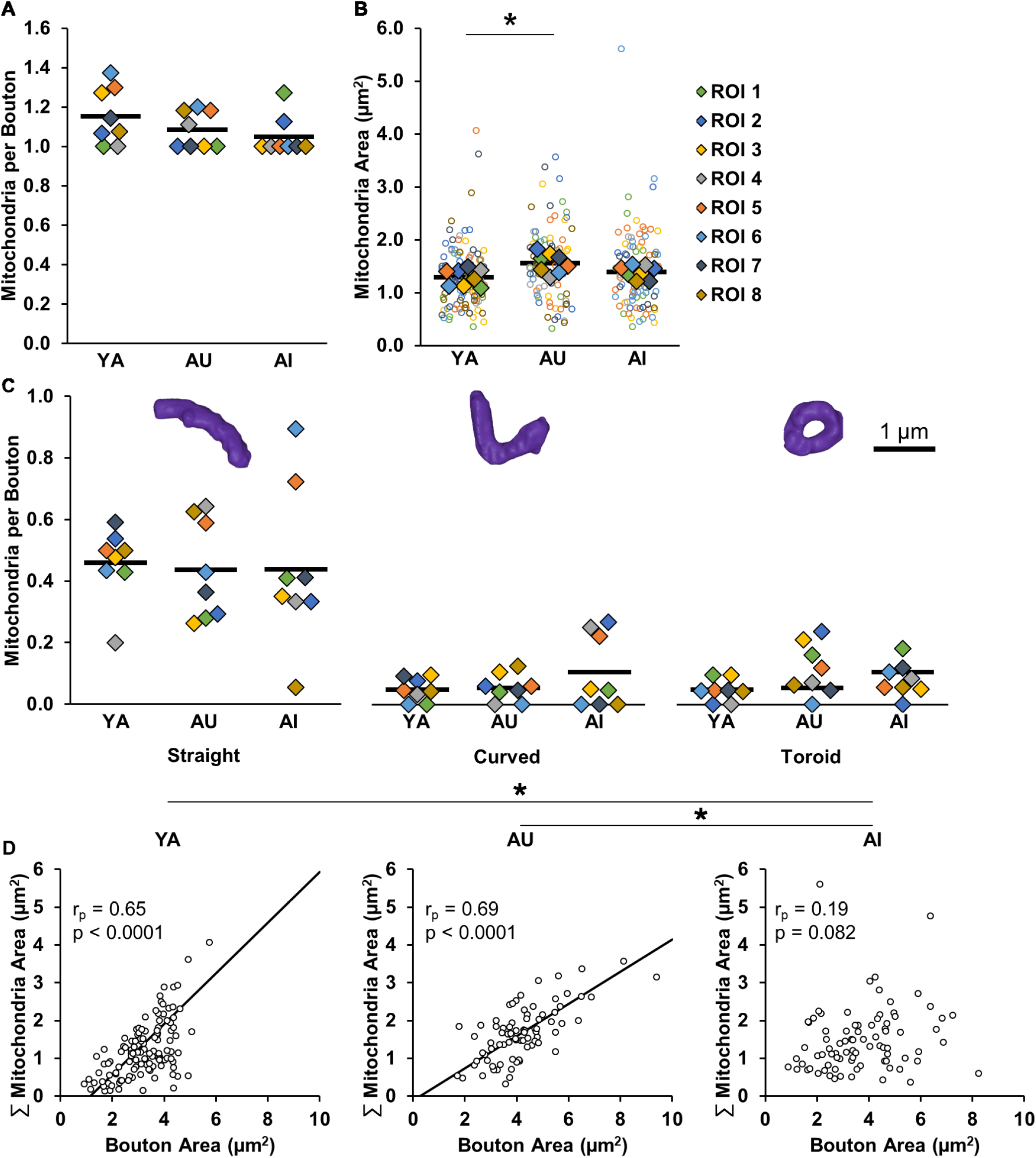
Presynaptic mitochondria. A) Mitochondrial density did not vary with age or cognitive status. B) Synaptic mitochondria from AU were larger than those in YA, while mitochondria from AI were statistically similar in size to both YA and AU. C) All animals had a greater proportion of mitochondria that were straight compared to both curved and toroidal. Within each morphology classification there were no differences between animals. D) While the sizes of mitochondria and boutons in YA and AU are highly correlated, these synaptic components are not correlated in AI. Open circles show each individual datapoint. Colored diamonds in A-C represent the average of measures taken in each of the ROIs, and horizontal black bars represent average across the ROIs. * p < 0.05.

Prior work demonstrates that dysmorphic mitochondria (e.g., toroids) are indicative of impaired energy production (Liu and Hajnóczky, 2011; Ahmad et al., 2013; Galloway and Yoon, 2013; Glancy et al., 2020). To test whether mitochondria morphology contributes to synaptic dysfunction in the AI marmoset, we quantified the number of mitochondria per bouton that were straight, curved, or toroidal in each animal. We found that while there were overall more straight mitochondria than curved or toroidal (Tukey’s HSD: straight vs curved: p = 9.56×10^-10^; straight vs toroidal: p = 9.56×10^-10^; curved vs toroidal: p = 0.95), there were no differences between animals (Figure 6B; two-way ANOVA: morphology F(1) = 72.94, p = 4.01×10^-17^; animal F(2) = 0.24, p = 0.79; interaction F(2) = 0.47, p = 0.75).

Since mitochondrial size correlates with energy production (Ivannikov et al., 2013; Ashaber et al., 2020), we next quantified mitochondria surface area to determine if smaller mitochondria are responsible for synaptic dysfunction in the AI marmoset. However, we found no differences between AI marmoset mitochondria and YA and AU mitochondria (Figure 6C; one-way ANOVA: F(2) = 5.84, p = 0.01; Tukey’s HSD: YA vs AU: p = 0.01, YA vs AI: p = 0.44, AU vs AI: p = 0.11).

Finally, we tested the hypothesis that mitochondria do not scale properly with bouton size. We found similarly strong correlations between these structures in YA and AU (Figure 6D; YA: r_p_ = 0.65, p < 0.0001, AU: r_p_ = 0.69, p < 0.0001; Fisher Z transformation: YA vs AU: p = 0.66). Remarkably, there was no correlation between bouton size and mitochondria size in AI (r_p_ = 0.19, p = 0.082; Fisher Z transformation: YA vs AI: p = 0.0002, AU vs AI: p < 0.0001). These results support our hypothesis that decorrelation between mitochondria and their boutons underlies synaptic dysfunction in AI.

## DISCUSSION

Like humans (Nyberg et al., 2020), marmosets exhibit heterogeneity in their cognitive aging profiles (Glavis-Bloom et al., 2022). Therefore, the marmoset is a powerful model for understanding the neural mechanisms that drive this variability and can help identify what causes some individuals to sustain age-related cognitive impairment, while others remain unaffected. In this study we examined the ultrastructural features of synaptic complexes and how they change as a function of age and age-related cognitive impairment. We used electron microscopy to visualize and quantify synaptic components in layer III of the dlPFC from one young adult and two aged marmosets. Each of the aged marmosets had previously undergone working memory assessment. One exhibited working memory function comparable to young adult marmosets, while an age-matched animal exhibited profound working memory impairment (Glavis-Bloom et al., 2022). While, consistent with prior studies (Peters et al., 2008; Dumitriu et al., 2010; Morrison and Baxter, 2012), we do see a loss of synapses with age, this loss was indistinguishable across the two aged animals. The main feature that did distinguish aging with and without cognitive impairment was a novel form of synaptic dysfunction: violation of the ultrastructural size principle. In particular, we found that while the sizes of presynaptic boutons were highly correlated with the sizes of synapse surface areas in a young adult and aged unimpaired animal, this correlation was significantly weaker in the age-matched impaired animal. Further, among boutons that contained mitochondria, the correlation between bouton size and mitochondrial size, which was strong in the young adult and aged unimpaired animal was entirely absent in the aged impaired animal. These findings are important in that they reveal a novel form of synaptic dysfunction that may contribute to age-related cognitive impairment. This study is, to our knowledge, the first to find evidence that age-related working memory impairments may stem from a violation of the ultrastructural size principle, and is the first characterization of synaptic ultrastructure in cognitively profiled marmosets.

### Aged marmosets have synapse loss consistent in severity to aged macaques

We focused here on the dlPFC because it is critical for working memory, and morphological and functional changes in this region have been observed early in the aging process (Morrison and Baxter, 2012; Upright and Baxter, 2021). Consistent with observations in macaques (Bertoni-Freddari et al., 2007; Peters et al., 2008;Dumitriu et al., 2010; Morrison and Baxter, 2012; Crimins et al., 2017), we find that aged marmosets have a marked (28%) reduction in synapse density compared to young. In addition, we find that the remaining synapses are larger in aged monkeys than young, and this is exacerbated with age-related working memory impairment. Prior work shows that the increased average synapse size is due to a selective loss of small synapses associated with thin spines that are critical for supporting working memory (Bertoni-Freddari et al.,2007; Dumitriu et al., 2010; Arnsten et al., 2012). To our knowledge, this is the first report of age-related synapse loss in the marmoset.

### Weaker correlation between the sizes of synapses and boutons with age-related cognitive impairment

Investigations of age-related changes to structural correlates of synaptic efficacy historically have focused on either presynaptic or postsynaptic analyses. The fact that pre and postsynaptic components are linearly related in size in young animals (i.e., aligning with the ultrastructural size principle) underscores the importance of investigating them together as a synaptic complex (Lisman and Harris, 1993; Pierce and Mendell, 1993; Pierce and Lewin, 1994). Importantly, to our knowledge, no study has investigated if the ultrastructural size principle is violated as a function of age or cognitive capacity. Therefore, we sought to understand perisynaptic alterations that could render synapses dysfunctional and may distinguish individuals that age without cognitive impairment from those that age with cognitive impairment. Consistent with prior work in aged macaques (Cork et al., 1990;Crimins et al., 2019), we found that aged marmoset presynaptic boutons were larger than young. Furthermore, congruent with the ultrastructural size principle and prior work in aged macaque dlPFC, there were strong linear relationships between synapse and bouton sizes in each of the marmosets. However, this correlation was significantly weaker in the aged marmoset with cognitive impairment compared to the aged unimpaired marmoset. Additionally, the ratio of synapse to bouton size was conserved between the young adult and aged unimpaired marmoset, but was significantly larger in the aged animal with working memory impairment, as compared to each of the other animals. This violation of the ultrastructural size principle indicates a form of synaptic dysfunction.

### Boutons containing mitochondria selectively account for violation of the ultrastructural size principle in age-related working memory impairment

The elevated firing rates that underlie working memory involve substantial synaptic activity. This activity is metabolically demanding, and the required energy is primarily supplied by synaptic mitochondria (Li et al.,2020). We thus hypothesized that a mismatch between energy supply and demand would be especially detrimental to maintenance of elevated firing, and is therefore a mechanism by which synaptic dysfunction occurs. To test this hypothesis, we divided synaptic complexes into two subgroups depending on whether the presynaptic bouton contained mitochondria or did not. We found that the increased ratio of synapse to bouton size in the AI marmoset was explained selectively by synaptic complexes with boutons that contained mitochondria. These findings lend support to the idea that mitochondria contribute to synaptic dysfunction underlying age-related working memory impairment, and leaves open the question of how this occurs.

### Evidence that mitochondria contribute to age-related working memory impairment through mismatched energy supply for synaptic demand

We proposed four explanations for how mitochondria could underlie synaptic dysfunction through mismatched energy supply for synaptic demand: 1) low mitochondrial density, 2) dysmorphic mitochondria, 3) reduced mitochondrial size, and 4) improper scaling. We tested each proposed mechanism by measuring structural correlates of mitochondrial energy production.

First, we hypothesized that the number of mitochondria per bouton is decreased in the AI marmoset compared to the YA and AU animals, and that the decreased mitochondrial density leaves synapses with insufficient energy to function properly. However, because mitochondrial density was equivalent across animals, lower density of synaptic mitochondria does not explain synaptic dysfunction in the AI marmoset. These results align with prior work reporting that young and aged macaques have similar mitochondrial density in dlPFC (Bertoni-Freddari et al., 2007; Hara et al., 2014), but differ from this prior work in that presynaptic mitochondrial density in macaque dlPFC correlated positively with working memory capacity (Hara et al., 2014). This positive correlation in macaques, however, was specific to mitochondria classified as healthy (i.e., straight), whereas density of mitochondria with other morphologies (e.g., curved, toroidal) were not predictive of memory capacity. Therefore, the health of the existing marmoset mitochondria, could explain mismatched energy supply for synaptic demand in age-related working memory impairment.

Mitochondrial morphology is indicative of their overall health, and thereby, ability to generate energy. Specifically, toroidal-shaped mitochondria produce less energy relative to that produced by straight mitochondria (Liu and Hajnóczky, 2011; Ahmad et al., 2013; Galloway and Yoon, 2013; Morrison and Baxter,2014; Glancy et al., 2020). Therefore, we hypothesized that an increased number of toroidal-shaped mitochondria in the AI compared to the YA and AU marmosets might lead to mismatched energy supply for synaptic demand and underlie synaptic dysfunction in AI. Similar to results from macaques (Hara et al., 2014), we found more straight mitochondria than curved and toroidal-shaped mitochondria in each marmoset, and no difference in the prevalence of toroidal-shaped mitochondria between young and aged animals. Further, both Hara et al. and our work do not find any correlation between the prevalence of toroidal-shaped mitochondria and performance on working memory tasks using delayed non-matching rules. Hara et al. did, however, find a negative correlation between toroidal-shaped mitochondria prevalence and performance on a delayed response task. This suggests that an association between mitochondrial morphology and working memory impairment may be task-dependent.

Mitochondrial size directly correlates with markers of synaptic efficacy (Pierce and Lewin, 1994; Ivannikov et al., 2013; Smith et al., 2016). Therefore, reduced mitochondrial size may lead to reduced energy production, resulting in mismatched synaptic energy supply and demand. However, our results do not support this as a mechanistic explanation for synaptic dysfunction because the average size of mitochondria in the aged animal with working memory impairment was equivalent to mitochondrial size in both of the other animals. However, the aged animal with conserved working memory had significantly larger mitochondria than the young animal. We hypothesized that the larger mitochondria reflect alignment with the ultrastructural size principle since the age-related increase in bouton size should be correlated with an increase in the size of presynaptic mitochondria. Further, the fact that mitochondria size did not vary between the aged impaired marmoset and either of the others suggests that synaptic dysfunction may be a result of improper scaling of mitochondria in the aged impaired animal.

Coordinated scaling of mitochondria with their presynaptic boutons is critical for maintaining effective synaptic transmission, especially during elevated firing that demands large amounts of energy. We assessed the degree to which synaptic mitochondrial size is directly proportional to the size of the presynaptic bouton, and found that the correlations were similarly strong between the young adult and the aged animal without cognitive impairment. Remarkably though, there was *no* correlation between mitochondria and presynaptic bouton size in the aged animal with working memory impairment, demonstrating violation of the ultrastructural size principle in age-related working memory impairment.

Therefore, we postulate that in the AU animal, working memory is maintained by effective synaptic transmission that is supported by matched energy supply and demand on a bouton by bouton basis. However, in the AI animal, working memory is impaired due to dysfunctional synaptic transmission caused by mismatched energy supply and demand stemming from a lack of coordination between the sizes of mitochondria and their containing boutons. An understanding of the signaling mechanisms that govern the coordination between synaptic components to maintain synaptic efficacy could lead to novel therapeutic targets to promote healthy cognitive aging.

### Summary

Here we have identified a novel mechanism by which age-related working memory impairment emerges: violation of the Ultrastructural Size Principle. We posit that this violation leads to synaptic dysfunction through mismatched energy supply and demand, disrupting the elevated firing necessary for working memory. To our knowledge, this work also represents the first examination of synaptic ultrastructure in the aged common marmoset. Finally, linking ultrastructure to cognitive capacity further validates the marmoset as a model for understanding the neural underpinnings of aging, and could reveal why aging itself is the biggest risk factor for neurodegenerative diseases.

## Acknowledgments

This research was supported by an AHA-Allen Initiative in Brain Health and Cognitive Impairment award made jointly through the American Heart Association and The Paul G. Allen Frontiers Group: 19PABH134610000AHA, a National Institutes of Health grant 1R21AG068967-01, Kavli Institute for Brain And Mind at UCSD (Innovative Research Grant 2021), grants from the Larry L. Hillblom Foundation and the Don and Lorraine Freeberg Foundation, and the Fiona and Sanjay Jha Chair in Neuroscience. We thank Katie Williams for assistance in the care of the marmosets and technical support.

## Conflict of interest statement

The authors declare no competing financial interests.

## Notes

### Competing Interest Statement

The authors have declared no competing interest.

